# Spatial protein analysis in developing tissues: a sampling-based image processing approach

**DOI:** 10.1101/163147

**Authors:** Karolis Leonavicius, Christophe Royer, Antonio Miranda, Richard Tyser, Anne-Marie Kip, Shankar Srinivas

## Abstract

Advances in fluorescence microscopy approaches have made it relatively easy to generate multi-dimensional image volumes and have highlighted the need for flexible image analysis tools for the extraction of quantitative information from such data. Here we demonstrate that by focusing on simplified feature-based nuclear segmentation and probabilistic cytoplasmic detection we can create a tool that is able to extract geometry based information from diverse mammalian tissue images. Our open-source image analysis platform, called ‘SilentMark’ can cope with noisy images and with crowded fields of cells to quantify signal intensity in different cellular compartments. Additionally, it provides tissue geometry related information, which allows one to quantify protein distribution with respect to marked regions of interest. The lightweight SilentMark algorithms have the advantage of not requiring multiple processors and graphics cards and can be run even with just several hundred MB of memory. This makes it possible to use the method as a web application, effectively eliminating setup hurdles and compatibility issues with operating systems. We test this platform on mouse pre-implantation embryos, embryonic stem cell derived embryoid bodies and mouse embryonic heart and relate protein localisation to tissue geometry.

## INTRODUCTION

One of the main areas of focus in developmental biology is how tissue complexity is generated and how diverse cell types are consistently organised into a complete organism. A common challenge in the field is the three-dimensional spatial analysis of protein distribution within tissues and cells (Lu-engo-Oroz et al., 2011; Schindelin et al., 2012). The function of a protein can vary depending on the cellular compartment it is localised to and differential expression of proteins across tissues during development is important in pattering those tissues. Cutting edge optical microscopy and image analysis tools are now routinely used for spatial analysis of developing tissues from various organisms (Barbier de Reuille et al., 2015; Chickarmane et al., 2010; Fernandez et al., 2010; Hodneland et al., 2013; Lou et al., 2014; Peng et al., 2010; Stegmaier et al., 2016). Given the variability of different tissues, existing software often have to be optimised to work for specific cases. For example, full cell segmentation tools like MorphographX (Barbier de Reuille et al., 2015) or Packing Analyzer (Aigouy et al., 2010) work extremely well with samples such as plants and Drosophila, where high quality fluorescence signals and low noise make it possible to segment complete cell shapes.

As the signal to noise ratio of the image drops or features become indistinct, it becomes more difficult to accurately segment and analyse individual cells. One of the most reliable solutions that is still widely used is manual segmentation (Trichas et al., 2011; Watanabe et al., 2014), because humans are adept at feature recognition. However, manual outlining is time consuming, which is particularly a problem for volume or time-lapse data. To automate image segmentation, there has been a shift towards machine learning segmentation approaches. A notable example is RACE (Stegmaier et al., 2016) which uses machine learning for tissue segmentation into individual cells. Arguably, this is one of the most successful approaches yet for automated segmentation, especially from membrane signals. However, it is still often impossible to correctly segment cells based on their outlines alone, because of limitations in image quality.

Dealing with multiple spatial and fluorescence variables raises additional challenges in terms of signal normalisation which is crucial in correcting for variations due to image acquisition or optical heterogeneity within tissues. A commonly used approach involves taking ratios of nearby cellular compartments, such as nuclei and cytoplasm. This has been previously used for example in investigating the Hippo pathway, where YAP localisation could be quantified by taking the ratio of nuclear and cytoplasmic fluorescence while at the same time normalising the raw signals (Halder et al., 2012).

Even in cases where image quality prevents reliable segmentation, the image data often contain a great deal of useful quantitative information. To extract such information, we have developed a novel sampling based approach to measure nuclear, cytoplasmic and plasma membrane fluorescence levels. Importantly, the localisation of these samples within the image volume is retained, which can then be used for detailed statistical analysis of the spatial distribution of the detected proteins. This approach works across the scale of individual cells to entire tissues. At the level of the cell, it allows us to quantify relative protein levels in different cellular compartments, while at the level of the tissue it allows us to identify cell populations based on relative protein expression, statistically test the difference between these populations and analyse how their location within the tissue correlates with protein levels.

Our approach uses well defined features such as nuclei to perform partial but robust segmentation using well established techniques (Hodneland et al., 2013; Illingworth and Kittler, 1987). Alternatively, well-resolved membrane signals or manually defined regions can also be used. If nuclei are used as points of reference, nearby voxels are used to sample nearby nuclear and cytoplasmic regions, identified based on nuclear stain intensity. Simultaneous sampling of different cell compartments allows us to normalize the data with respect to experimental variation and tissue optical heterogeneity. Regions of membrane sampling are located by using a combination of detected nuclei positions and intensity of a membrane stain. Membrane signal quantification can be further improved by using commercially available software such as Imaris to manually outline membrane regions, which are then automatically partitioned by our software into contacting (basolateral), exposed (apical) and junctional domains. Since the spatial distribution of proteins is of particular interest to developmental biologists, we have also incorporated the ability to define points or regions of interest in the image volume, so that the relative positions of these sampled regions with respect to these points of interest can be recovered.

We have implemented this approach as a software package called ‘SilentMark’, designed for general use. The package is a standalone GUI-based application written in Matlab. The algorithm has also been implemented in Python as a web application (http://data.dropletgenomics.com). Some limitations of our sampling based approach are that it does not provide information on cell shape, the total sum of fluorescence or facilitate cell tracking. However, the unique aspect is that it allows one to process routine lower quality images, investigate multiple cell compartments and quantify the spatial distribution of fluorescence within cells and tissues. Here we demonstrate the use of our algorithm on mouse pre-and post-implantation embryos, as well as mouse embryonic stem cell derived embryoid bodies.

## RESULTS AND DISCUSSION

To quantify protein levels in different cellular compartments and determine the relative proportion of the plasma membrane in different domains (apical, junctional, basolateral), we developed a sampling based algorithm named ‘SilentMark’, implemented in Matlab and Python. The overall strategy relies on detecting well resolved tissue elements, such as nuclei or membranes and then analysing their closest environment to take samples of subcellular region fluorescence. Depending on which cell marker is available, the first step is to detect well resolved objects such as nuclei or membranes, after which their closest environment is analysed.

### Nucleus based fluorescence sampling

Available nuclear detection techniques differ in their accuracy, throughput and noise tolerance. One of the fastest and most robust methods for spherical object detection is based on Hough transform (Illingworth and Kittler, 1987), which is a tensor voting technique for the most probable circle positions within an image. The original algorithm is developed for 2D image analysis. However it can be extended for 3D measurements by slicing the image into sections, analysing them as separate images and then deconvolving into 3D objects, as done previously (Stegmaier et al., 2016). In this case, confocal image stacks were analysed as separate images to detect circular nuclear cross-sections, the coordinates of which were then merged to generate a 3D point cloud of nuclear centers of mass (Figure 1 A). Using these nuclear coordinates as reference points, a region from the confocal image is cropped around each detected point and classified according to the intensity of the nuclear stain as either a cytoplasmic or nuclear region. The fluorescence from both regions is then averaged with cytoplasm pixels being weighed according to their distance from the detection point (Figures 1 B and 4 A, B). This was done to represent the probability that pixels closer to the nucleus are more likely to belong to the same cell as the nucleus. The fluorescence levels along with the nucle-ar coordinates were recorded and this formed the basis of segmentation and quantification.

**Figure 1:**
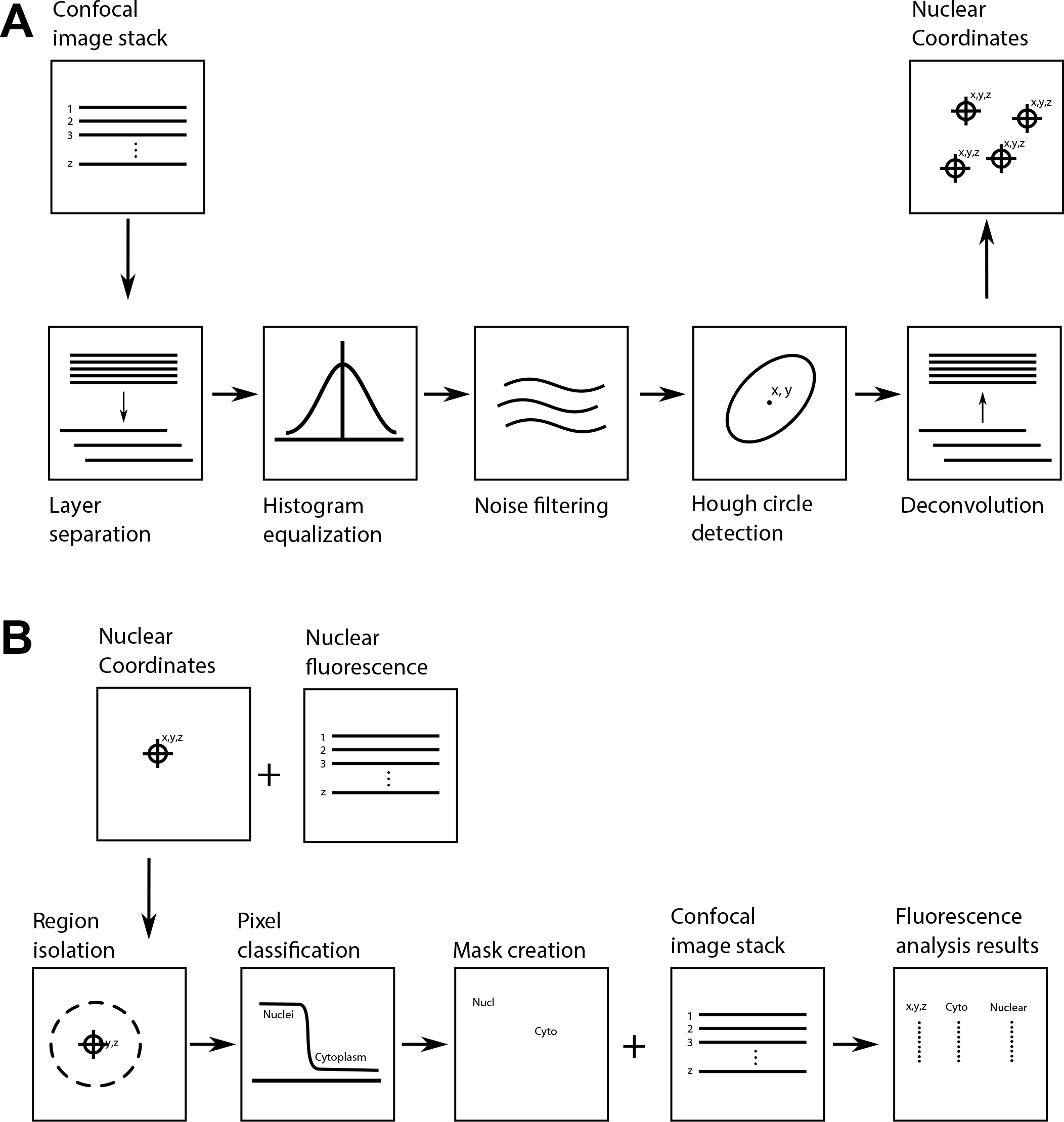
Schematic illustrating strategy for detecting the coordinates of the nuclear center of mass (A) and for extracting nuclear and cytoplasmic fluorescence based on nuclear coordinates (B).

If a membrane stain was available in addition to the nuclear stain, it was possible to add another fluorescence reading for exposed and contacting membranes regions, which is particularly useful for looking at membrane localised proteins. To detect outside surface, the membrane fluorescence marker was thresholded and the outer pixels were assigned to the closest nucleus as an exposed membrane sample. To detect the contacting membrane samples, a line was drawn between two neighbouring nuclei and the highest intensity region for membrane fluorescence along the line was designated as a contacting membrane sample and was assigned to both nuclei.

### Membrane based fluorescence sampling

In cases where nuclear markers are not available as a reference, membrane fluorescence, if available, can be used to create the points of reference for subsequent analysis. To detect membrane regions in space, a 3D stack was processed using contrast limited adaptive histogram normalization (CLAHE) and Gaussian blur to reduce noise. Following this, a 3D Canny Edge detector (Bähnisch et al., 2009) was applied to find membrane edges, to which spheres were fitted using an outlier tolerant RANSAC algorithm. The detected membranes were represented by a location of the fitted sphere and a vector representing membrane curvature and direction (Figure 2 A). This information would then be the starting point for sampling membrane and cytoplasmic fluorescence intensities. The advantage of this method is that it looks for fragments of membranes, which is more tolerant to noise than other approaches that rely on total membrane segmentation. These detected membrane regions can now be used as sampling masks for membrane and cytoplasmic fluorescence. The mask considers a narrow region in space around the detected point to sample membrane fluorescence and draws a normal to membrane surface to sample cytoplasm on the either side. These masks are used on 3D image stacks to analyse membrane and cytoplasmic fluorescence distribution in space (Figure 2 B). Membrane signal quantification can be improved by using commercially available software Imaris to manually outline membrane regions, which SilentMark then automatically partitions into contacting, exposed and junctional domains (Figure 4 D), where proteins are likely to play different roles.

**Figure 2:**
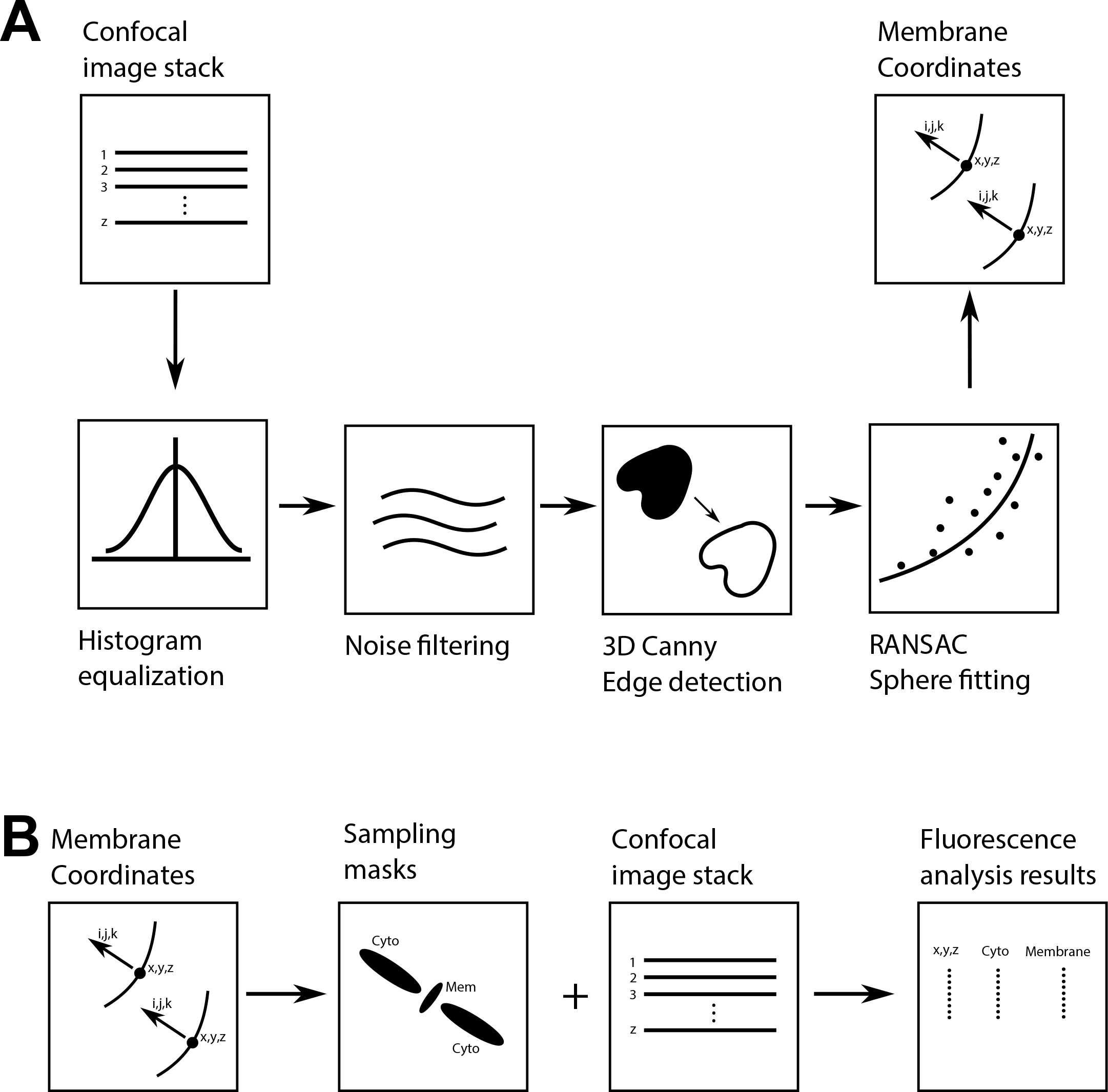
Schematic illustrating the membrane fragment detection algorithm (A) and analysis of membrane and cytoplasmic fluorescence based on membrane fragment detection (B).

### Development of ‘SilentMark’ software

We have developed these sampling based algorithms into a software package designed for general use. The package is a standalone application, based on a GUI designed on Matlab. The algorithm has also been implemented in Python as a web application (http://data.dropletgenomics.com). It accepts 3D images as stacks of tiff files or the Zeiss confocal microscopy ‘.lsm’ format. The user is required to enter two intuitive and robust parameters – an estimate of the diameter of the nuclei and an estimate of nuclear channel brightness for thresholding. These two minimal parameters allow one to work with different types of tissues and image qualities. All the remaining parameters needed for the analysis are derived and optimised by the software.

The software will output a list (csv format) of detected objects (nuclei or membrane fragments), each of which will have a measure of nuclear and cytoplasmic fluorescence as well as membrane fluorescence, if it is labeled. The data will also contain object coordinates to describe spatial protein distribution, automatically calculated distance to exposed surface and distance to regions of interest, if they are designated manually in 3D. This information can then be processed with statistical software tools for further analysis.

### Method validation with CAD images

To validate this probabilistic sampling based approach, we first used computer generated 3D images to compare the accuracy of our automated sampling based approaches against standard manual outlining (using Imaris) (Figure 3). Mockup tissues composed of several spherical objects representing cells were created on Google Sketch-Up and converted to a stack of images, which were then filled with pixels of known brightness. Unsurprisingly, manual outlining was the most accurate method of image segmentation and in addition to quantifying fluorescence intensity, could also be used to estimate the proportion of ‘exposed’ cell surface, that is, cell surface not in contact with surrounding cells. In general, intensity measurement errors were below 10%, except for feature detection relying on membrane fragments. The most likely source of error for membrane based feature detection came from the detection algorithm while fitting spheres, which is responsible for identifying both membrane segment position and orientation.

**Figure 3:**
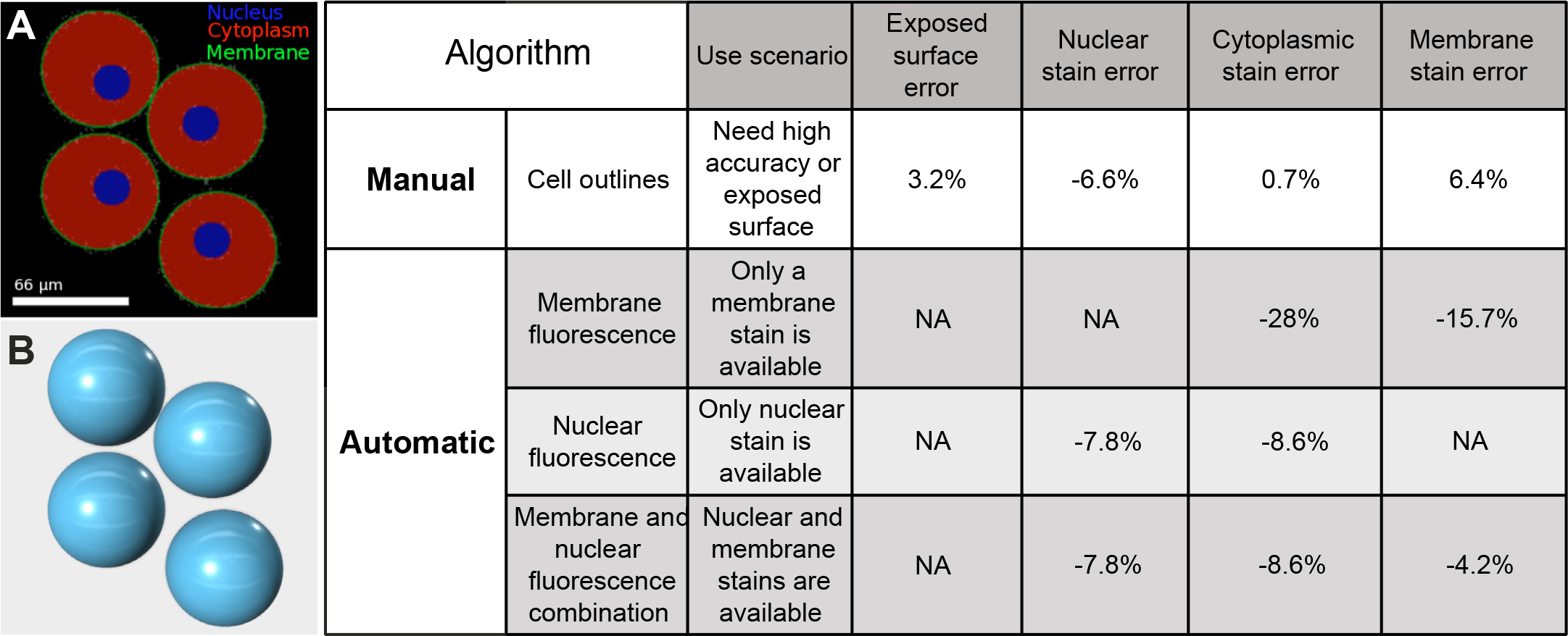
Accuracy of sampling based image segmentation. A) A section of a computer generated image for testing image analysis methods. B) A 3D rendering of computer generated objects for testing image analysis methods. Method accuracy was assessed by comparing measured fluorescence levels to known intensity values in computer generated images. Four models each consisting of four to six cells were used in this test.

### Method validation with pre-implantation embryos

The mouse pre-implantation embryo is a model system where cell fate is strongly influenced by tissue geometry (Sasaki, 2010; Wennekamp et al., 2013) The underlying basis for cell type specification in the pre-implantation embryo has been studied in great depth and this large amount of existing information makes it ideal for validating our analytical approach.

During the process of the first cell fate decision in mammals, the protein YAP shuttles between the cytoplasm (where it is inactive) and the nucleus (active), in a tightly regulated manner. The differential localisation of YAP protein in outside and inside cells ultimately determines cell fate in this context and the ratio of nuclear to cytoplasmic YAP reflects the position of cells. In outside cells, YAP is in the nucleus where it can interact with nuclear TEAD4 to drive the expression of tissue specific genes such as Cdx2 to give rise to the trophectoderm. In inside cells, YAP is excluded from the nucleus and these cells go on to become the pluripotent inner cell mass (Nishioka et al., 2009; Niwa et al., 2005; Sasaki, 2017).

To estimate the errors resulting from the sampling based approach we used manual cell outlining to create a ground-truth dataset of pre-implantation embryos stained for YAP (n = 67 embryos, comprising 20 2-cell, 15 8-cell, 17 16-cell, 12 32-cell and, three 64-cell embryos. Examples in Figure 5 and supplementary movie 1). For nuclear and cytoplasmic fluorescence measurements mean error was lower than 5% (Figure 4 E), which indicates a low bias during automated sampling.

**Figure 4:**
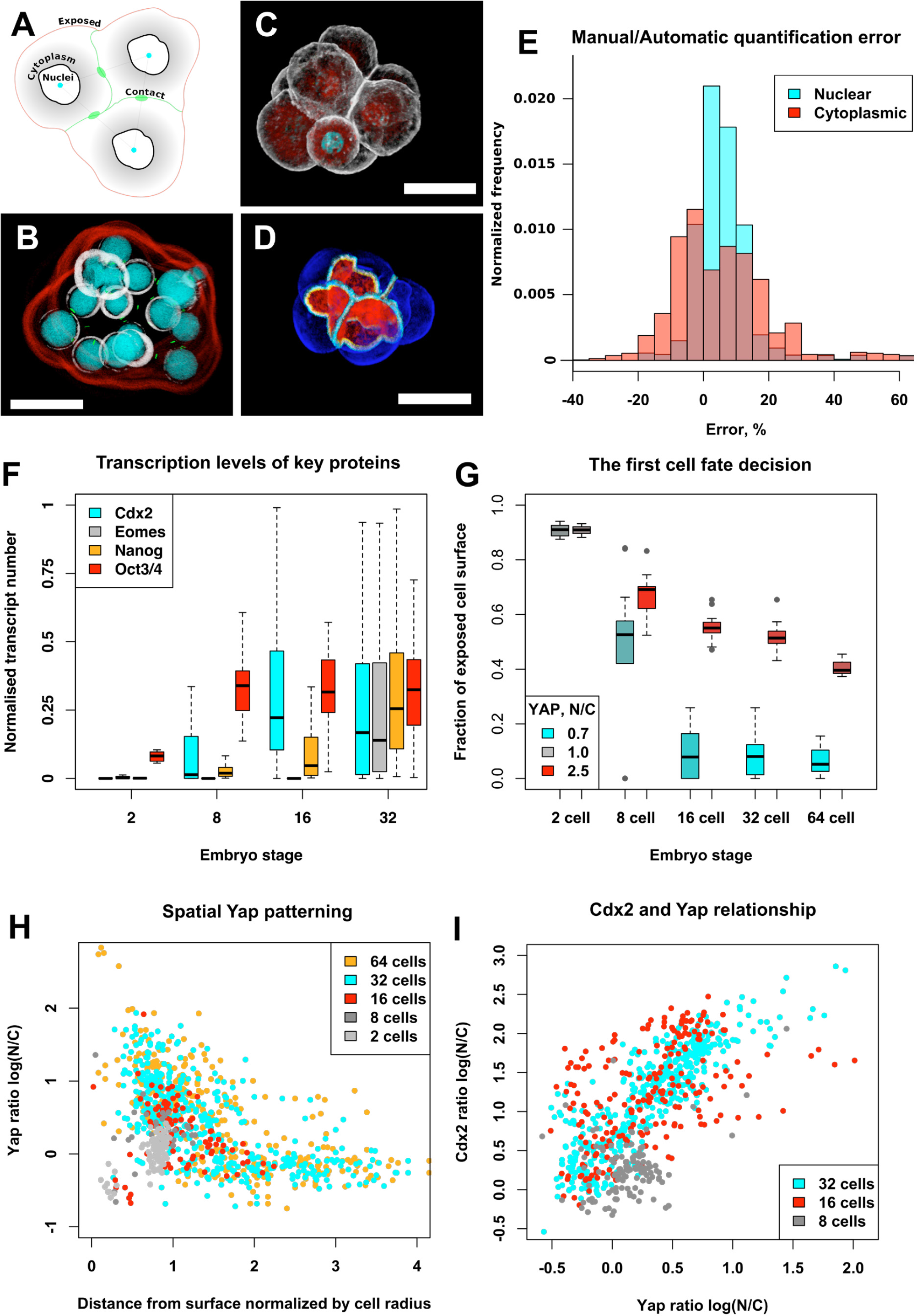
Quantifying lineage detreminants during mouse pre-implantation development. A) sampling algorithm for detecting three sub-cellular regions; B) 3D rendering of different sampling regions. Red and green are exposed and contacting membranes respectively; gray is cytoplasm and; cyan is nucleus. C) 3D rendering of a mouse embryo (gray – actin, red – YAP, cyan – DAPI); D) 8-cell embryo in which cell membranes have been manually segmented and then membrane regions automatically sub-segmented into ‘exposed’ (blue), ‘junctional’ (white) and basolateral (red). E) Error distribution of automated sampling based quantitation compared with manual segmentation based quantitation. F) Levels of key transcription factors during development. G) YAP distribution in the early pre-implantation embryo. H) Spatial YAP patterning during pre-implantation development. I) YAP and CDX2 relationship in the morula and early blastocyst. Scale bars in all panels are 50µm.

We used our automated sampling based approach to explore the relationship between YAP and CDX2 levels and the extent to which this correlates with a blastomere’s position, as measured by exposure to the outside or distance of a cell from the surface. We used embryos stained for YAP, CDX2 and the DAPI. Published single cell RNA-seq data on mouse pre-implantation embryos (Deng et al., 2014) (GEO accession: GSE45719) indicates that Cdx2 becomes upregulated at the 8 cell stage (Figure 4 F, n = four 2-cell, four 8-cell, four 16-cell and three 32-cell embryos). Our analysis shows that at the 8-cell stage, blastomeres begin to show signs of segregating into two populations, with high or low nuclear to cytoplasmic YAP ratios (Figure 4 G and example in Figure 5. N = 20 2-cell, 15 8-cell, 17 16-cell, 12 32-cell and three 64-cell embryos). More robust segregation of these two populations occurs at the 16-cell stage, approximately 12 hours after Cdx2 transcript appearance at the 8-cell stage. Analytical detail makes it possible to demonstrate that populations are separated gradually, as YAP translocates to the cytoplasm in inside cells (Figure 4 H). It is also evident, that the CDX2 – YAP correlation is stronger at the 32-cell stage than the 16-cell stage (Figure 4 I and example in Figure 5. N = seven 8-cell, nine 16-cell and six 32-cell embryos). This is consistent with the presence of additional stabilizing mechanisms for robust lineage specification (Rayon et al., 2014; Ralston et al., 2010).

**Figure 5:**
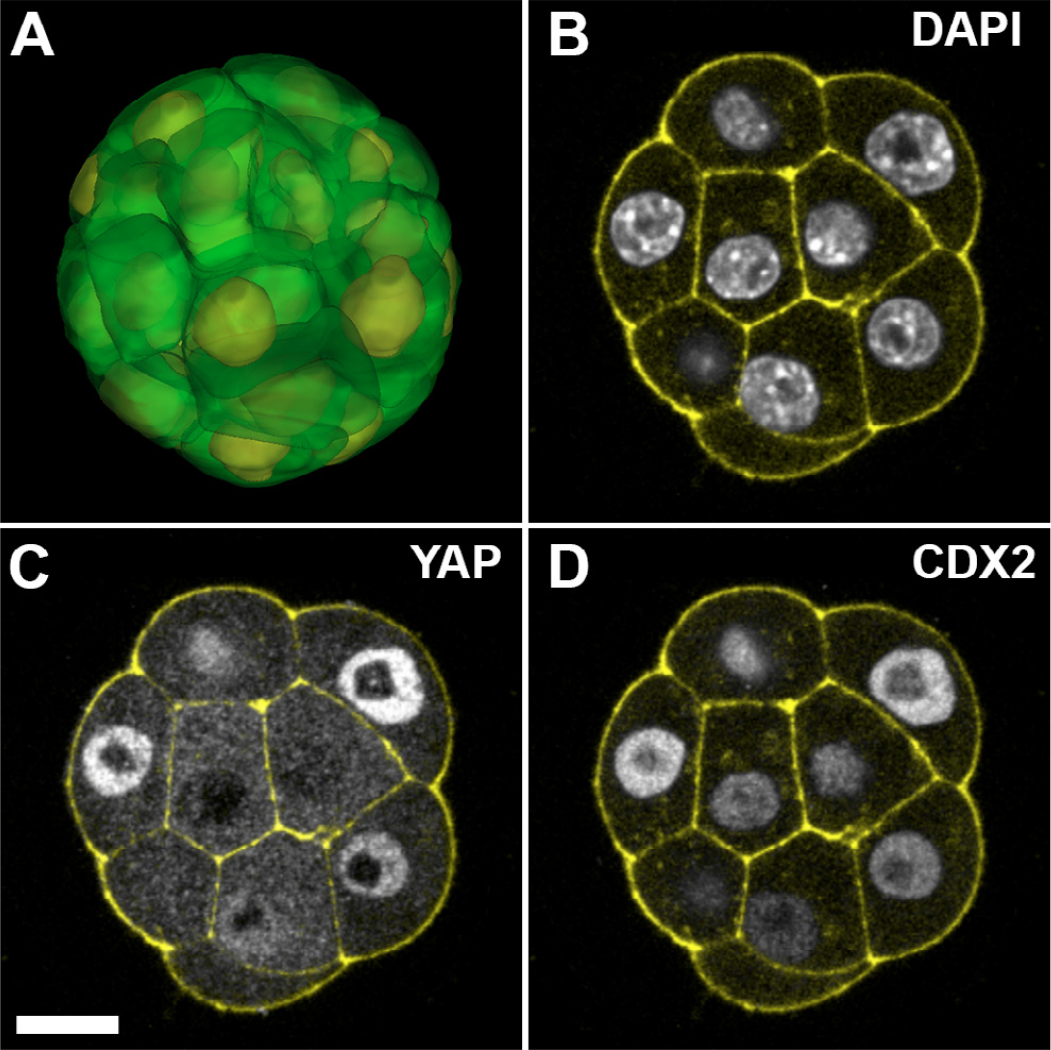
A) an example of a manually segmented 16-cell mouse embryo, with cell outlines in green and nuclei in yellow. B, C and D) Example of a 16-cell mouse embryo stained for DAPI, YAP and CDX2. In all images cell outlines are in yellow, stained with Phalloidin (f-actin). The scale bar is 50 µm.

### Method validation with ES cell derived embryoid bodies

Similar to mouse pre-implantation embryos, mouse embryonic stem cell aggregates also display inside–outside patterning during embryoid body (EB) formation in vitro (Doetschman et al., 1985). The outer layer of cells forms an endoderm like layer around an inner core that is epiblast like. These aggregates tend to be composed of crowded cells (Figure 6 A, B) and represent a more challenging task for both manual and automated segmentation or sampling approaches. To test its capabilities, we applied our sampling based image approach to investigate endoderm formation during EB differentiation. We stained EB for the endoderm marker GATA6 (Figure 6 B) that specifically stains outside cells and quantified the relative levels of nuclear GATA6 expression with respect to the surface of EB (n = 6). Quantitative statistical analysis (Figure 6 C) shows preferential GATA6 to the outer layer of the embryoid body (distance to outer surface is smaller than two cell radii).

**Figure 6:**
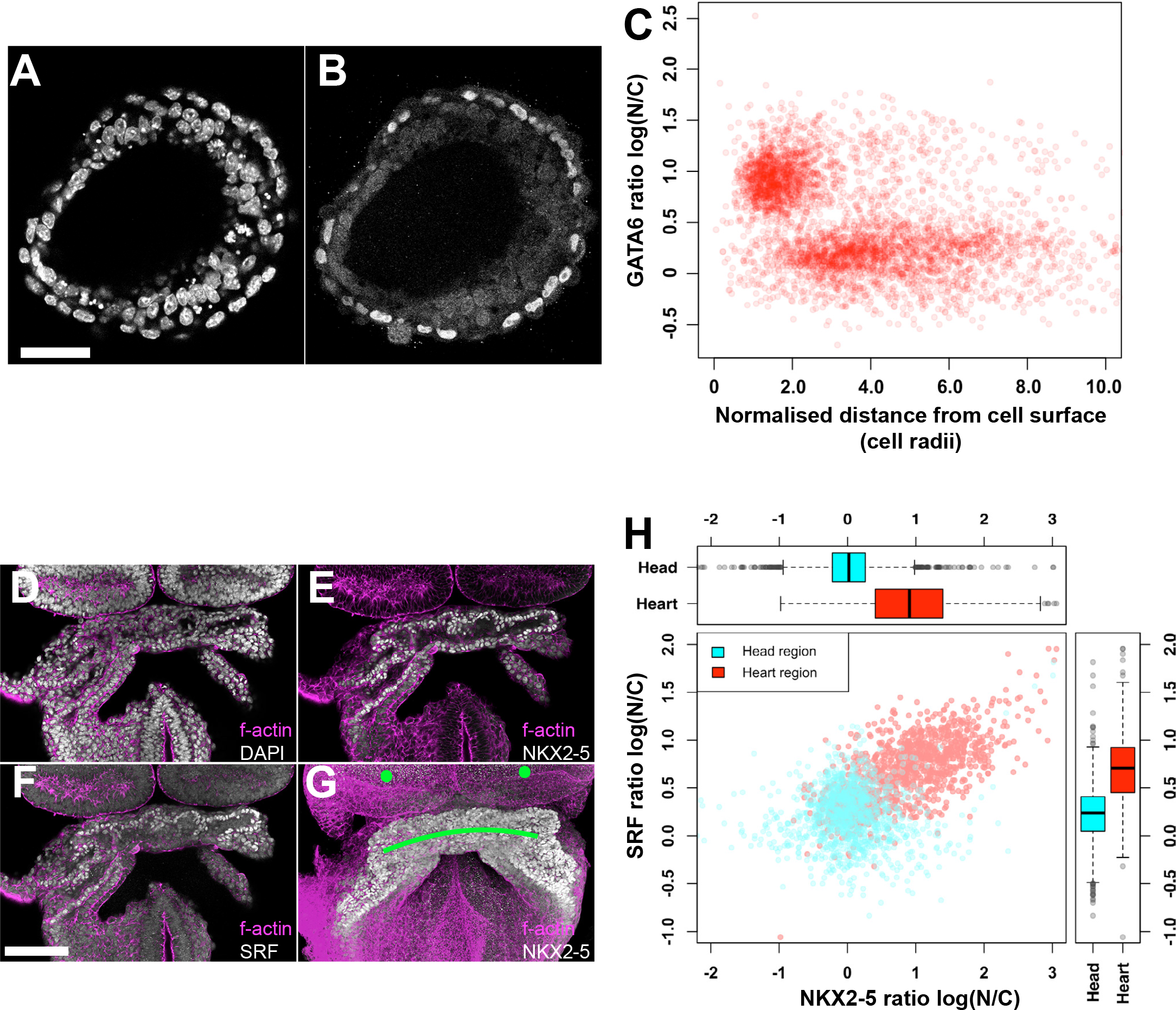
Automated quantitative analysis of differentiated mouse stem cell derived embryoid bodies and mouse embryonic heart. A and B) Optical sections of a differentiating mouse stem cell embryoid body stained with DAPI (A, cell nuclei) and GATA6 (B, marking outer endoderm layer). C) GATA6 localization in the embryoid body as a function of distance from exposed surface (n = 6 EBs). The approximately singe cell wide outer layer is distinct from the remaining embryoid body. D - F) Optical section of an embryonic heart stained for F-actin (magenta in all panels, showing cell outlines), DAPI (D, grey, all cell nuclei), NKX2-5 (E, grey nuclei in cardiac crescent) and SRF (grey nuclei in cardiac crescent). G) Maximum intensity projection of image volume of embryonic heart, with F-actin in magenta and NKX2-5 in grey. The green dots and line are regions of interest marking the head folds and cardiac crescent respectively. H) Localization and correlation of SRF and NKX2.5 transcription factors, with boxplots comparing proteins in different embryo regions. The blue and red dots were clustered by the Clara algorithm based on SRF and NKX2-5 levels and their distance from the heart and head folds. p-values are <0.001 (t-test). Scale bar in panel A represents 50 µm and in panel F 100µm.

### Method validation with embryonic hearts

While mouse pre-implantation embryos present an ideal case for testing automatic image segmentation, because it is possible to also manually outline the cells for comparison, they do not necessarily represent a typical mammalian tissue. A more representative mammalian tissue is the embryonic heart (Figure 6 D-G) with a larg-er number of cells and closely spaced nuclei.

To test the ability of our image analysis approach for this problem, we examined the localisation of NKX2-5 and SRF, two transcription factors important for cardiomyocyte differentiation (Schlesinger et al., 2011) in 8.0 days post coitum embryos (Figure 6 E, F). In order to represent protein levels of these two transcription factors we used a ra-tio of nuclear to cytoplasmic fluorescence, as the latter is expected to represent background signal for internal normalisation. To reduce the dynamic range of the measurements and normalise the distribution of the values, we used a natural logarithm of the ratio. In addition to these normalised fluorescence measurements, we also measured cell distance to two regions, the head folds and cardiac crescent, manually designated using our software (green dots and line in Figure 6 G).

The four variables formed a dataset which was clustered using partitioning around medoids (PAM, R ‘Clara’ package) (van der Laan et al., 2002). The method was chosen over other clustering approaches (Wiwie et al., 2015) due to computational efficiency in dealing with large datasets. The analysis verified as expected that both proteins were preferentially expressed in the heart region and there was a region-specific correlation between nuclear levels of NKX2-5 and SRF (Figure 6 H).

In summary, we have presented a sampling based 3D image analysis approach, designed for relative quantification of protein levels in microscopy images of complex tissues. Our approach is the first to simultaneously quantify nuclear, membrane and cytoplasmic fluorescence, while also marking regions of interest to extract spatial information relating to the quantified signals. Automation of analyses enabled by our program will enable high-throughput quantification and statistical analysis of spatial protein organization. As 3D microscopy is versatile and widespread, we expect that our publicly available open-source software will be useful not only for developmental biology but also more broadly, in the context of cell biology.

## METHODS

### Mouse strains, husbandry and embryo collection

All animal experiments were carried out according to UK Home Office project license PPL 30/3155 and 30/2887 compliant with the UK animals (Scientific Procedures) Act 1986 and approved by the local Biological Services Ethical Review Process. All mice were maintained in a 12-hour light-dark cycle. Noon of the day finding a vaginal plug was designated 0.5 dpc (days post coitum). To obtain embryos, C57BL/6 males were crossed with CD1 females (Charles River). Embryos of the appropriate stage were dissected in M2 medium (Sigma-Aldrich) at room temperature. Pre-implantation embryos were collected by oviduct and uterus flushing.

### Embryonic stem cell culture

ES cells were cultured in T-25 flasks coated with 0.1% w/v. gelatin and used at passage nr. 50-60. Cell culture media consisted of 90% DMEM medium (Gibco), 10% fetal calf serum (FCIII, Hyclone), 1mM sodium pyruvate, 1mM non-essential amino acid mix, 0.1mM beta-mercaptoethanol, 2mM L-glutamine, and 1000U/mL leukemia inducible factor (LIF). Cells were passaged every 2 days and seeded at ~400,000 cells per T25 flask. Embryoid bodies were formed by aggregating cells in hanging drops and culturing for 2 days before fixation.

### Antibodies

Vectashield with DAPI (H-1200) was purchased from Vector laboratories, Phalloidin atto647 was purchased from Sigma (65906-10MMOL). Mouse anti-YAP monoclonal (sc-101199) and goat anti-Nkx2.5(N-19) (sc-8697) polyclonal antibodies were purchased from Santa Cruz biotech. Rabbit anti-CDX2 (#3977S) monoclonal and rabbit anti-pERM(T567) (#3149P) monoclonal antibodies were purchased from Cell Signaling. Rat an-ti-Uvomorulin/E-cadherin (U3254-100UL) monoclonal antibody was purchased from Sigma. Rabbit anti-SRF (PA5-27307) polyclonal antibody was purchased from Thermo Fischer. We used the following secondary antibodies from Life Technologies: goat anti-rabbit AlexaFluor 488 (A11008), donkey anti-goat AlexaFluor 488 (A11055), donkey anti-mouse AlexaFluor 555 (A31570), donkey anti-rabbit AlexaFluor 647 (A31573).

### Immunofluorescence on pre-implantation embryos and embryoid bodies

Embryos and EB were fixed for 20 minutes at room temperature in 4% PFA. Samples were then washed 3 times in PBT-0.1% for 10 min, permeabilized in PBT-0.25% for 40 min and washed again in PBT-0.1%. The tissues were transferred to a blocking solution for 1h at 4 oC. Primary antibodies were then added to the solution and incubated overnight at 4 oC. The embryos were washed in PBT-0.1% and incubated for 1h at 4 oC in PBT-0.1% with the secondary antibodies, then subsequently washed 2 times in PBT-0.1% for 15 min and mounted in Vectashield with DAPI at least 6 hours prior to imaging.

### Immunofluorescence on post-implantation embryos

Dissected embryos were fixed for 1 hour at room temperature in 4% PFA. The embryos were then washed 3 times in PBT-0.1% for 15 minutes, permeabilized in PBT-0.25% for 40 minures and washed again 3 times in PBT-0.1%. The embryos were transferred to blocking solution overnight at 4 oC. Primary antibodies were then added to the solution and incubated at 4 oC. The embryos were washed 3 times in PBT-0.1% and incubated overnight at 4 oC in PBT-0.1% with the secondary antibodies, then subsequently washed 3times in PBT-0.1% for 15 minutes and mounted in Vectashield with DAPI at least 24 hours prior to imaging.

### Software and statistics

The software and graphical user interface were written on MATLAB (v. 2015b, Mathworks) and compiled using the in-built compiler for Mac OSX and Windows operating systems. The algorithm has also been implemented in Python as a web application (hosted by Droplet Genomics at http://data.dropletgenomics.com). Source code for Silent Mark is available upon request. 3D image visualization was done using Volocity software (Perkin Elmer) and Imaris software (v 6.1) was used for manual cell outlining and analysis. R statistics package alongside ‘cluster’ library was used for automatic normalized cluster detection (PAM algorithm) and cell population analysis.

## AUTHOR CONTRIBUTIONS

K.L. developed the analysis method in discussion with S.S. K.L., C.R. and A.K. performed early embryo and stem cell experiments.

R.T. performed the embryonic heart experiments. K.L. and A.M designed statistical anal-yses. S.S, K.L. and C.R. wrote the manuscript.

## ACKNOWLEDGEMENTS

K.L. was supported by a Biotechnology and Biological Sciences Research Council (BBSRC) Doctoral Training Program grant to the University of Oxford (BB/J014427/1). This work was supported through a BBSRC equipment grant (BB/F011512/1) and Wellcome Senior Investigator Award to S.S. (Ref. 103788/Z/14/Z).

